# Baraitser–Winter Syndrome Hotspot Mutation R196H in Cytoskeletal β–actin Reduces F–actin Stability and Perturbs Interaction with the Arp2/3 Complex

**DOI:** 10.1101/2024.03.20.585892

**Authors:** Johannes N. Greve, Dietmar J. Manstein

## Abstract

Baraitser–Winter cerebrofrontofacial syndrome (BWCFF) is the most common and best–defined clinical entity associated with heterozygous single–point missense mutations in cytoskeletal β–actin. Patients present with distinct craniofacial anomalies and neurodevelopmental disabilities of variable severity. To date, the most frequently observed variants affect residue R196 of cytoskeletal β–actin, with the variant p.R196H being the most common. Patients carrying the p.R196H variant are likely to suffer from pachygyria, probably due to neuronal migration defects contributing to the development of abnormally thick convolutions of the cerebral cortex. Here, we describe the recombinant production, purification and biochemical characterization of the BWCFF hotspot variant p.R196H. The stability and nucleotide interaction of monomeric p.R196H are unaffected, indicating a disease mechanism involving incorporation of p.R196H protomers into actin filaments. Incorporation of the variant strongly affects F–actin stability and polymerization dynamics, consistent with the position of residue R196 close to the helical axis of the actin filament and an important interstrand contact. The changes observed include an increased critical concentration of polymerization, a reduced elongation rate and an increase in the rate of filament depolymerization. In the Arp2/3–generated branch junction complex, which is essential for cell migration and endocytosis, R196 is located at the interface between the first protomer of the nucleated daughter filament and the Arp2 subunit of the Arp2/3 complex. Assays probing the interaction of p.R196H filaments with the Arp2/3 complex show a reduced efficiency of branch generation. Branch stability is impaired, as evidenced by a reduction in the number of branches and spontaneous debranching events. Furthermore, in their interaction with different types of cytoskeletal myosin motors, p.R196H filaments show isoform–specific differences. While p.R196H filaments move WT–like on lawns of surface–immobilized non–muscle myosin–2A, motility on myosin–5A is 30 % faster.

## Introduction

Cytoplasmic actin filaments are composed of the ubiquitous actin isoforms cytoskeletal β– and γ–actin, encoded by *ACTB* and *ACTG1* (1). Essential cellular processes like cell migration, adhesion, division, and signal transduction depend on an intact actin cytoskeleton and a robust remodeling of cytoskeletal actin structures, which enables cells to respond to intra– and extracellular stimuli by conferring cytoskeletal plasticity (2–4). Actin remodeling, involving assembly and disassembly of actin filaments and higher order F–actin structures is tightly regulated by a large number of G– and F–actin binding proteins. This regulation allows spatio–temporal control of cellular actin dynamics and drives functional compartmentalization of the actin cytoskeleton (5,6).

Heterozygous missense mutations in *ACTB* or *ACTG1* result in a broad spectrum of non–muscle actinopathy phenotypes observed in affected patients (7–11). To date, the best described clinical entities associated with missense variants in *ACTB* and *ACTG1* are non–syndromic hearing loss, due to missense mutations in *ACTG1* (12–14), and the Baraitser–Winter cerebrofrontofacial syndrome (BWCFF) caused by missense variants in *ACTB* and *ACTG1* (7,8,15). BWCFF is associated with brain malformations and neuronal migration defects that lead to neurodevelopmental disorders and intellectual disabilities. The patients present with distinct craniofacial features such as facial malformations, microcephaly, ptosis, hypertelorism, and ocular coloboma syndrome (7,8). The more severe cases of BWCFF are regularly observed in patients carrying variants in *ACTB* (10). Missense variants associated with BWCFF are scattered throughout the *ACTB* and *ACTG1* sequences, with no apparent preference for any particular region. Nonetheless, specific variants are more often observed in affected patients and are thus termed hotspot variants. These include the variant p.R196H in cytoskeletal β–actin, which is by far the most frequently observed variant among all BWCFF variants and causes a prototypical BWCFF phenotype (8). Two additional variants were observed at this position, p.R196S and p.R196C, both resulting in a similar phenotype. Patients carrying these variants present with BWCFF–specific craniofacial malformations, muscular hypotonia, moderate to severe intellectual disability and delayed speech (7,8).

Due to the large number of possible disease mechanisms, the link between the observed clinical phenotype in the patient and the perturbation of the molecular mechanisms of actin dynamics caused by the mutant actin remains a major challenge in the field of actin-associated diseases. Postulated disease mechanisms include functional haploinsufficiency due to instability and rapid degradation of the mutated actin (16), distorted regulation of cellular actin dynamics due to direct or allosteric perturbances of actin–actin and actin–ABP interaction interfaces (17–20), and formation of toxic rod–like actin aggregates (21).

Here, we report the biochemical characterization of the BWCFF hotspot variant p.R196H in cytoskeletal β–actin, the variant with the most reported carriers to date. Residue R196 is located in a flexible loop in subdomain 4 of the actin protomer, away from the sensitive hinge region, the nucleotide–binding cleft and the target–binding cleft, which forms a major binding site for monomer–binding ABPs (**Figure 1A**).

**Figure 1:**
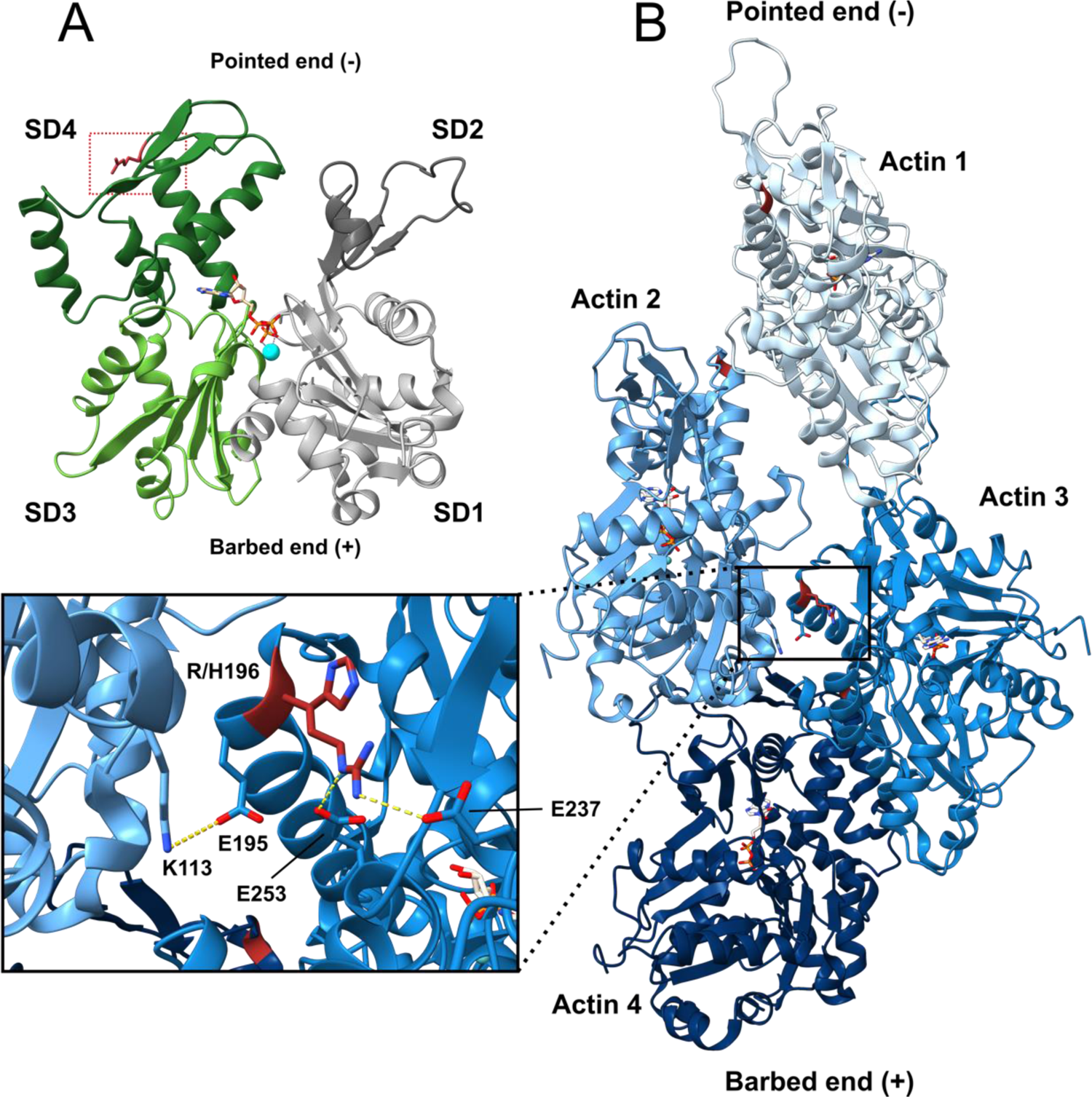
Location of the mutated residue R196 in the G– and the F–actin structure. **(A)** Structural model of cytoskeletal β–actin (based on PDB–ID: 2BTF (58)). Subdomains (SDs) of actin are colored in grey (SD1), dark grey (SD2), light green (SD3) and green (SD4). ATP is bound to the nucleotide binding site, the coordinated Mg^2+^ ion is depicted in cyan. The mutated residue R196 is located in SD4 and colored red. **(B)** Structural model of the human β–actin filament in the Mg^2+^–ADP bound state (PDB–ID: 8DNH (59)). Actin protomers are shown in different shades of blue. ADP is colored in light grey, the residue R196 is colored red. The zoom–in shows the position of residue R196 and the variant residue H196 close to the longitudinal axis of the filament. Salt–bridges established between R196 and E237, E253 in the same actin protomer are indicated by yellow dotted lines. The salt-bridge formed by the neighboring residue E195 and residue K113 in the protomer of the opposing filament strand is shown by the yellow dotted line (H–bond cut–off: 4Å).

After integration of the monomer into a filament, R196 is found close to the helical axis of the actin filament. Here, the neighboring residue E195 is involved in the formation of a salt–bridge with K113 in the opposite protomer (**Figure 1B**). This interaction constitutes one of the few lateral contacts across the filament axis that are involved in filament stabilization. These lateral contacts are far fewer in number than the more extensive longitudinal contacts formed between the molecules in the individual filament strands (22). Furthermore, R196 forms several electrostatic interactions with charged residues of the same actin protomer (**Figure 1B**).

We show that mutation R196H strongly perturbs actin assembly dynamics *in vitro* by increasing the critical concentration for actin polymerization, reducing the rate of filament elongation and increasing the rate of filament depolymerization. With respect to the interaction with key actin–binding proteins, we show that p.R196H interacts less efficiently with the Arp2/3 complex and demonstrate myosin– isoform specific changes in the ability of p.R196H filaments to function as myosin motor tracks.

## Results

### Production of cytoskeletal β–actin and the variant p.R196H

We produced cytoskeletal β–actin and the p.R196H variant as an actin–thymosin–β4 fusion protein (23) in the baculovirus/Sf9 insect cell expression system. To facilitate purification and prevent polymerization during recombinant production and initial purification steps, the monomer sequestering protein thymosin–β4 carrying a C–terminal His_8_–tag was fused to the β–actin C-terminus (**Figure 2A**). Immobilized metal affinity chromatography was employed for the initial purification of the fusion protein, followed by chymotrypsin–mediated cleavage to remove the C–terminal linker and thymosin– β4 moiety. This process yields recombinant cytoskeletal actin devoid of any non–native residues and with correct N–terminal acetylation of aspartate–2 (24,25). The recombinant actin was purified further by cycles of polymerization and depolymerization. A typical preparation started with 2×10^9^ cells and yielded 6–8 mg of purified protein for WT cytoskeletal β–actin and 3–5 mg of purified protein for the p.R196H variant (N=3 preparations for WT β–actin and N=4 preparations for p.R196H). The lower yields obtained with the p.R196H variant can be attributed to a reduced ability to form filaments during the final purification steps comprising polymerization and depolymerization cycles.

**Figure 2:**
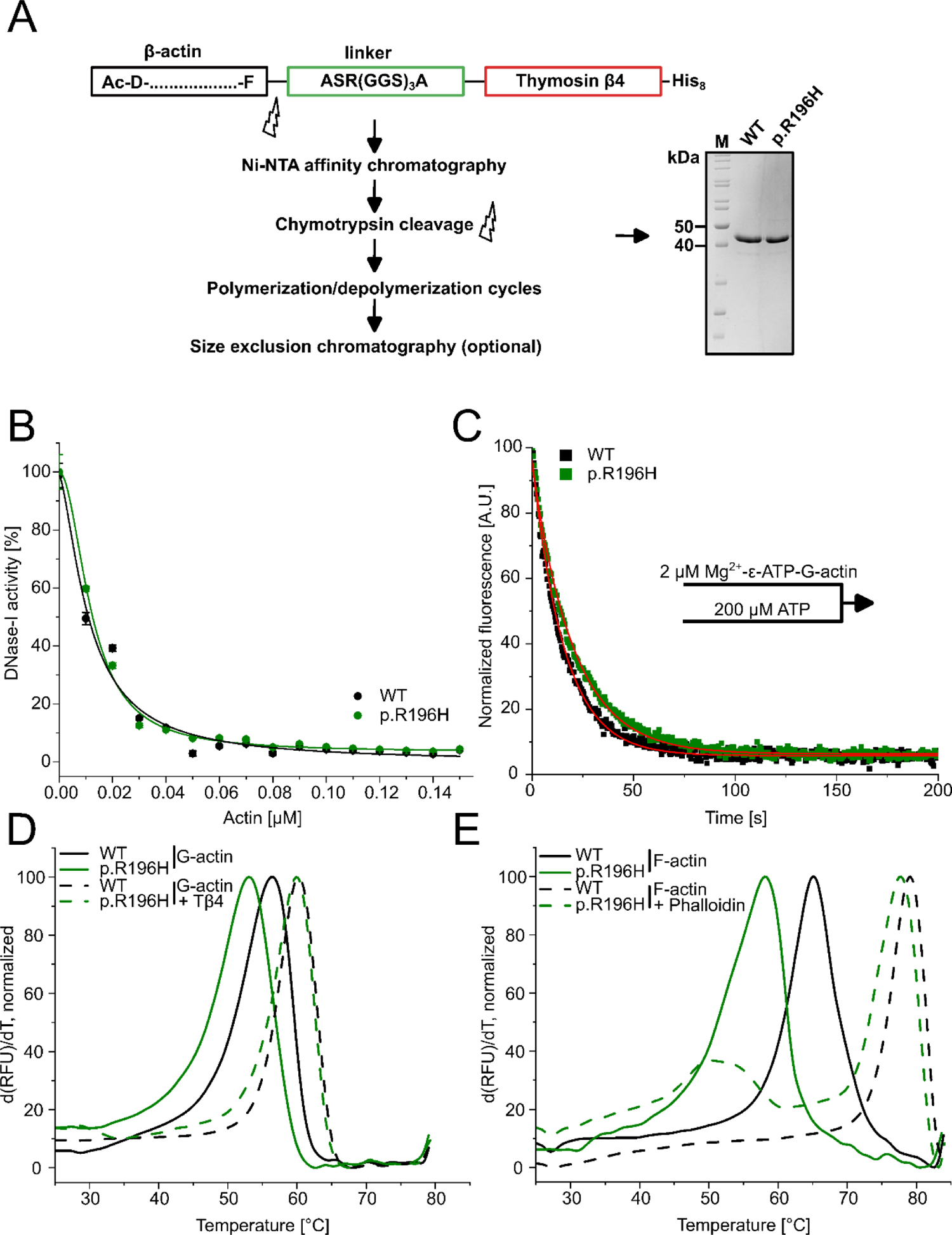
Analysis of actin folding, nucleotide exchange and thermal stability. **(A)** Schematic representation of the strategy used to purify recombinant human cytoskeletal β–actin WT and p.R196H. The SDS–gel shows purified WT and p.R196H cytoskeletal β–actin. **(B)** Inhibition of DNase–I activity by monomeric p.R196H and β–actin WT. Data is the mean of three individual experiments ± SD. A Hill equation was fitted to the data, which yields the half–maximal inhibitory concentration (IC_50_). **(C)** The rate of nucleotide dissociation (*k*_-T_) was determined for monomeric p.R196H and β–actin WT using fluorescently labeled ATP (ε–ATP). Shown are representative experimental traces to which single–exponential functions (red) were fitted. **(D, E)** The protein denaturation temperature (T_M_) of monomeric (D) and filamentous (E) p.R196H and WT actin were determined by DSF. Representative experimental traces are shown. The thermal denaturation temperature was derived from the peak of the melting curves.

### Effect of mutation R196H on protein folding and thermal stability

The effect of the R196H mutation on the folding of the protein was first assessed using a DNase–I inhibition assay (**Figure 2B**). Misfolded or denatured actin shows a reduced capacity of DNase–I inhibition, as previously shown (26). We observed no significant change in the observed IC_50_–value of p.R196H compared to WT actin (IC_50_ (WT) = 10.7 ± 1.1 nM; IC_50_ (p.R196H) = 12.4 ± 0.6 nM).

Efficient nucleotide binding is important for the full functionality and stability of G–actin. We investigated the ability of monomeric p.R196H and WT actin to interact with ATP by performing nucleotide dissociation experiments with fluorescent ε–ATP using a stopped–flow apparatus (**Figure 2C**). To determine the dissociation rate *k*_-T_, we monitored the displacement of fluorescent ε– ATP from the nucleotide binding site after mixing with excess unlabeled ATP. For p.R196H monomers (*k*_-T_ = 0.051 ± 0.003 s^-1^), we observed a 29% decrease in *k*_-T_ compared to WT monomers (*k*_-T_ = 0.065 ± 0.002 s^-1^).

Using differential scanning fluorimetry (27), we analysed the thermal denaturation temperature (T_M_) of the monomeric (G–) form and filamentous (F–) forms of WT and p.R196H actin (**Figure 2D, E)**. The R196H mutation results in a slight decrease of approximately 3 °C in the T_M_ for mutant G–actin (T_M_ = 53.1 ± 0.1 °C) compared to the WT protein (T_M_ = 56.3 ± 1.1 °C). The actin–sequestering protein thymosin–β4 binds a significant amount of G-actin *in vivo* and it has previously been shown that binding of thymosin–β4 to G–actin significantly increases T_M_ compared to uncomplexed G–actin (28). Thus, we performed thermal denaturation experiments in the presence of thymosin–β4. We found that the presence of thymosin–β4 increased T_M_ for both WT (60.3 ± 0.1 °C) and p.R196H actin (52.4 ± 0.6 °C), offsetting to a large extent the observed difference in T_M_ between uncomplexed WT and p.R196H G–actin (**Figure 2D**). Next, we determined the T_M_ of WT and p.R196H filaments. In accordance with previous studies (29), we found that compared to G–actin, WT filaments show an increase in T_M_ of approximately 8.5 °C to 64.7 ± 0.5 °C. Filaments containing p.R196H show a smaller increase in T_M_ of 5.8 °C from 53.1 ± 0.1 to 58.9 ± 0.6 °C (**Figure 2E**). Their more asymmetric peak shape, which partially overlaps with the melting curve obtained for monomeric p.R196H, is consistent with a disturbed G– to F–actin ratio. Differential scanning fluorimetry experiments with WT and p.R196H F–actin in the presence of the *Amanita phalloides* toxin phalloidin confirmed this finding (**Figure 2E**). Phalloidin is commonly used in biochemical studies to produce stable filaments at concentrations below the critical concentration required for polymerization (≥ 0.1 µM). It binds F-actin with high affinity and stabilizes the filamentous form by increasing the number of lateral and longitudinal contacts and restricting the relative movement of the two strands of the F–actin helix (30).The stabilizing effect of phalloidin on WT filaments results in an increased T_M_ in the presence of phalloidin (T_M_ = 78.7 ± 0.5 °C) and an apparent reduction in the full width at half maximum (FWHM). We interpret this as an increase in cooperativity of the denaturation process due to reduced structural flexibility of the actin filament. In the case of p.R196H filaments, two peaks were observed in the presence of phalloidin. A larger peak at approximately the same temperature as in the WT samples (T_M_ = 76.9 ± 0.5 °C), and a second smaller peak with a maximum at approximately 52 °C, which matches the melting curve of uncomplexed p.R196H G–actin. Thus, it appears that phalloidin is unable to completely shift the G– to F–actin ratio in the p.R196H actin sample to the F–form, stabilizing p.R196H F–actin less efficiently than WT F–actin.

### Mutation R196H results in perturbed polymerization dynamics and rapid filament depolymerization

We analysed the effect of the R196H mutation on the polymerization capacity of actin by performing single–filament polymerization studies using fluorescently labeled WT or p.R196H actin and total internal reflection fluorescence microscopy (TIRFM) (**Figure 3A, B**). At a concentration of 1 µM, we observed efficient nucleation and elongation of WT filaments resulting in an average barbed–end elongation rate of 14.9 ± 0.4 nm s^-1^. We rarely found filaments in experiments performed with p.R196H at the same concentration. Most of the observed p.R196H filaments were loosely attached to the surface, while one end showed slow elongation (3.1 ± 0.6 nm s^-1^), apparently interrupted by periods of depolymerization (**Figure 3D**). To increase the number of filaments observed, we increased the concentration of actin to 2 µM. Indeed, at 2 µM we found significantly more p.R196H filaments in all experiments performed. The filaments showed an average elongation rate of 9.0 ± 0.2 nm s^-1^, which is significantly lower than the elongation rate of WT filaments at 1 µM (14.9 ± 0.4 nm s^-1^) and at 2 µM (23.3 ± 0.7 nm s^-1^) (**Figure 3B**). Since BWCFF is caused by heterozygous mutations, WT actin in patient cells produced by the unaffected allele may attenuate the observed molecular defects if it is capable of copolymerizing with p.R196H actin. Therefore, we repeated the TIRFM–based experiments with 1:1 mixtures of p.R196H and WT actin. Experiments performed with a final concentration of 1 µM actin (0.5 µM p.R196H, 0.5 µM WT) produced sufficient filaments to determine the rate of elongation. Filaments in these experiments showed an average elongation rate of 7.3 ± 0.1 nm s^-1^. This rate is significantly faster than the elongation rate of 0.5 µM pure WT actin (4.8 ± 0.2 nm s^-1^) and significantly slower than the elongation rate of 1 µM pure WT actin (14.9 ± 0.4 nm s^-1^). Therefore, p.R196H copolymerizes with WT actin into heterofilaments containing mutant and WT protein. The presence of WT actin attenuates the polymerization defect of p.R196H, but p.R196H still significantly affects the elongation rate of the heterofilaments. Increasing the actin concentration to 2 µM (1 µM p.R196H, 1 µM WT) also increased the observed elongation rate of the heterofilaments to 14.9 ± 0.2 nm s^-1^, which is not significantly different to the elongation rate of 1 µM pure WT actin, but still slower than the observed elongation rate of 2 µM pure WT actin (23.3 ± 0.7 nm s^-1^).

**Figure 3:**
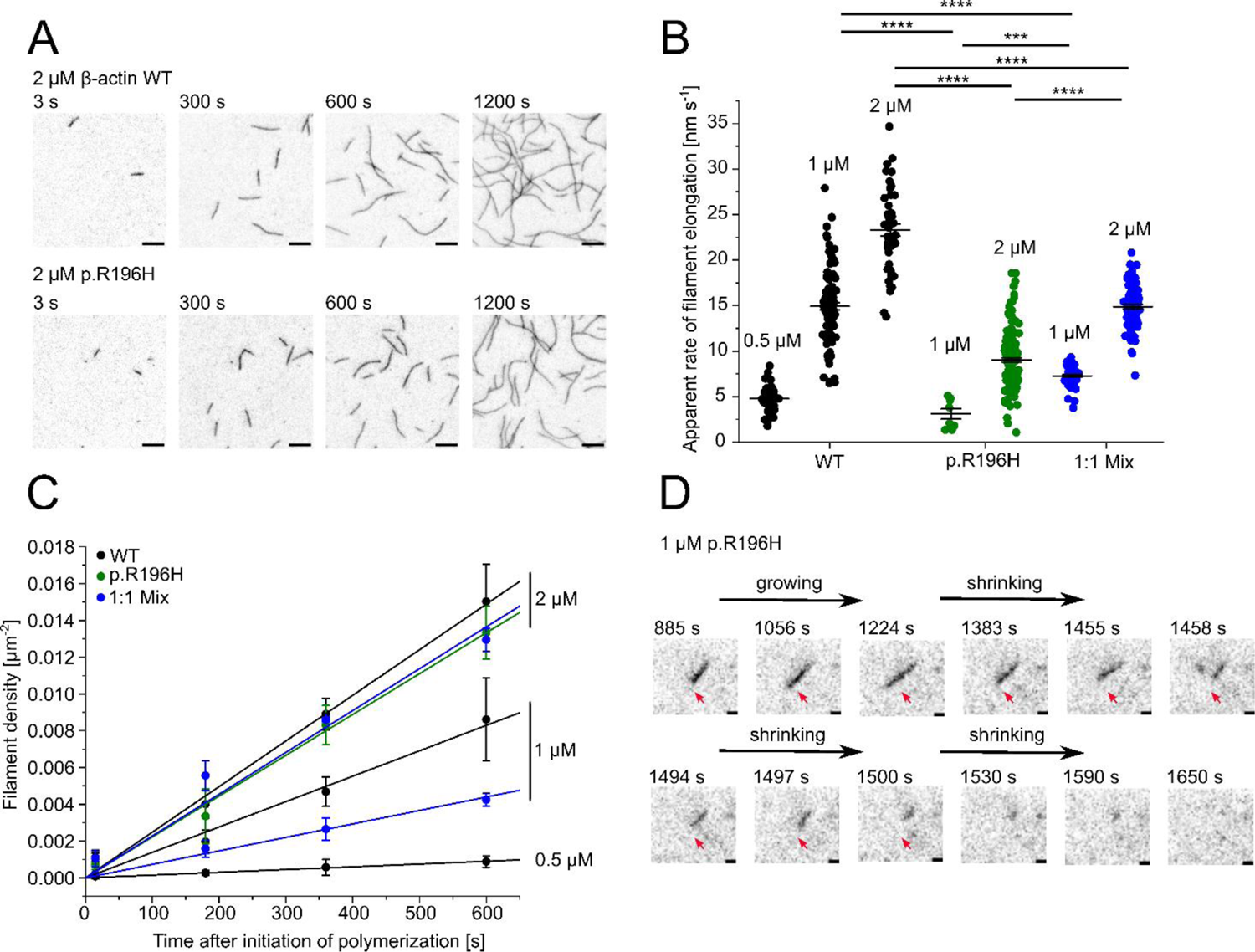
Analysis of the polymerization capacity of WT and p.R196H β–actin at different actin concentrations by TIRF microscopy. **(A)** Polymerization of p.R196H and WT β–actin (0.5 µM, 1 µM and 2 µM; 10% ATTO–655 labeled) was induced by salt–shift and the progression of the reaction was tracked by TIRF microscopy. Shown are representative micrographs at the indicated points in time from experiments performed with 2 µM p.R196H and WT actin. Scale bar corresponds to 10 µm. **(B)** The elongation rates of individual filaments were determined by manual tracking of the elongating barbed ends of the filaments. Every data point represents an individual filament (WT (0.5 µM) = 55 filaments, WT (1 µM) = 97 filaments, WT (2 µM) = 47 filaments, p.R196H (1 µM) = 8 filaments, p.R196H (2 µM) = 159 filaments, Mix (1 µM) = 61 filaments, Mix (2 µM) = 90 filaments). Data is shown as the mean ± SEM. **(C)** Nucleation efficiencies were calculated by determining the filament density [µm^-2^] over the time of the polymerization experiments. A linear regression was applied to the data to determine the apparent nucleation efficiency (*k*_nuc_) (Table 1) Data is shown as the mean ± SD of 3–4 individual experiments **(D)** Exemplary p.R196H filament in polymerization experiment performed with 1 µM p.R196H actin. The filament is loosely attached to the surface. One end of the filament (red arrow) shows slow polymerization followed by depolymerization. Scale bar corresponds to 1 µm.

**Table 1.**
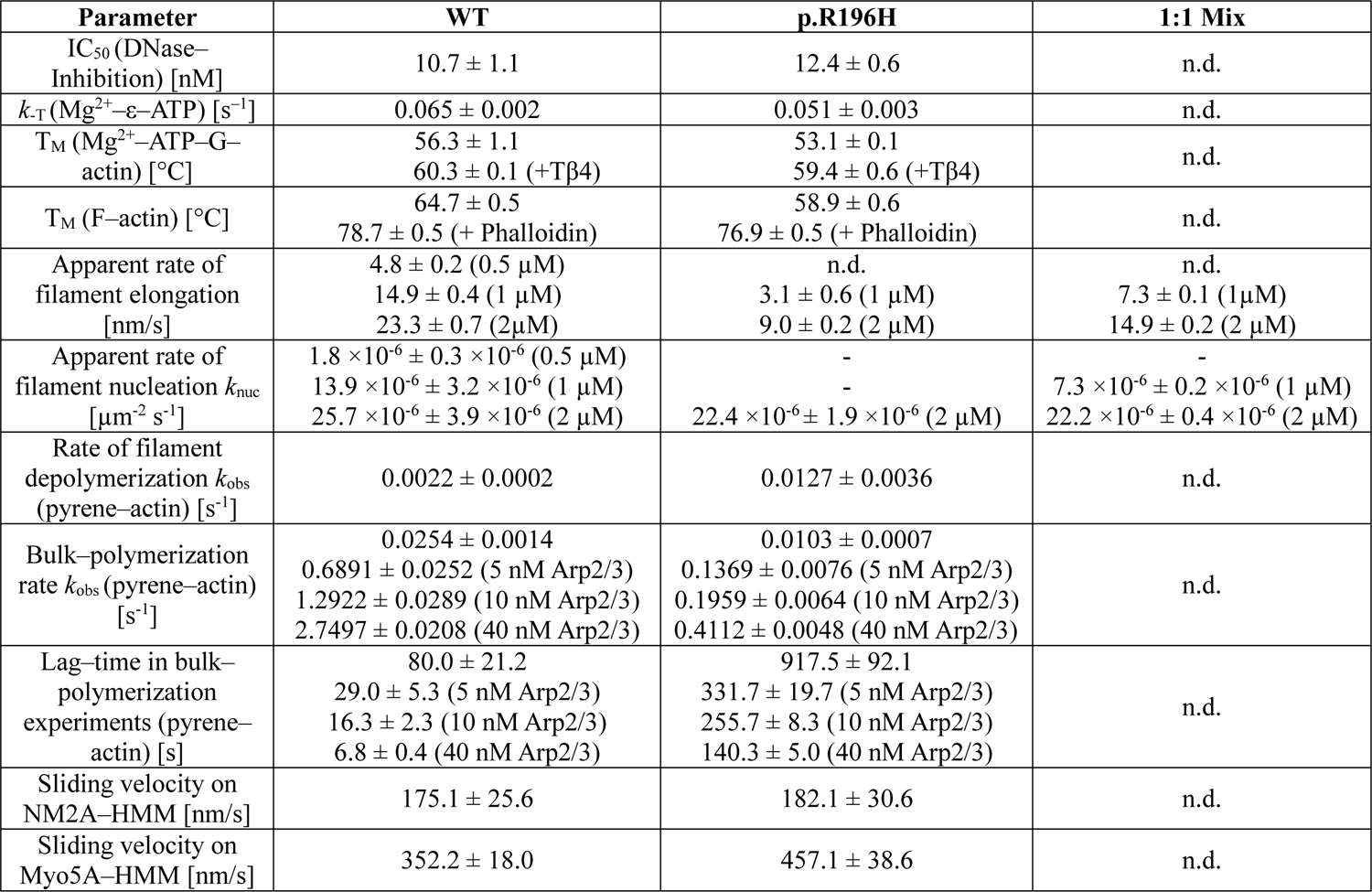
Comparison of biochemical parameters, velocities and rates measured in experiments with human cytoskeletal β–actin and variant p.R196H.

Every observed filament in the TIRFM–based experiments is the result of a successful nucleation event. Therefore, we determined the filament density in WT and p.R196H experiments at distinct points in time to quantify the nucleation efficiency of WT and p.R196H actin at different concentrations (**Figure 3C**). We observed a clear increase in the rate of filament nucleation (*k*_nuc_) of pure WT actin with increasing concentration (**Table 1**, **Figure 3C**). We compared the nucleation efficiency of WT, p.R196H and the 1:1 mix of both at a final concentration of 2 µM actin and found only minimal differences at this concentration (**Table 1**). As previously stated, we rarely found p.R196H filaments at a concentration of 1 µM. These results point to a significant defect of filament nucleation at concentrations below 2 µM.

To investigate the polymerization defect further, we performed sedimentation experiments with increasing concentrations of pre–polymerized WT and p.R196H F–actin (**Figure 4A**). We found less p.R196H actin at every concentration in the pellet fraction compared to WT actin. Applying a linear regression to the data revealed that the critical concentration of p.R196H actin (c_c_ = 0.66 ± 0.08 µM) is increased 5.5–fold compared to WT actin (c_c_ = 0.12 ± 0.06 µM).

**Figure 4:**
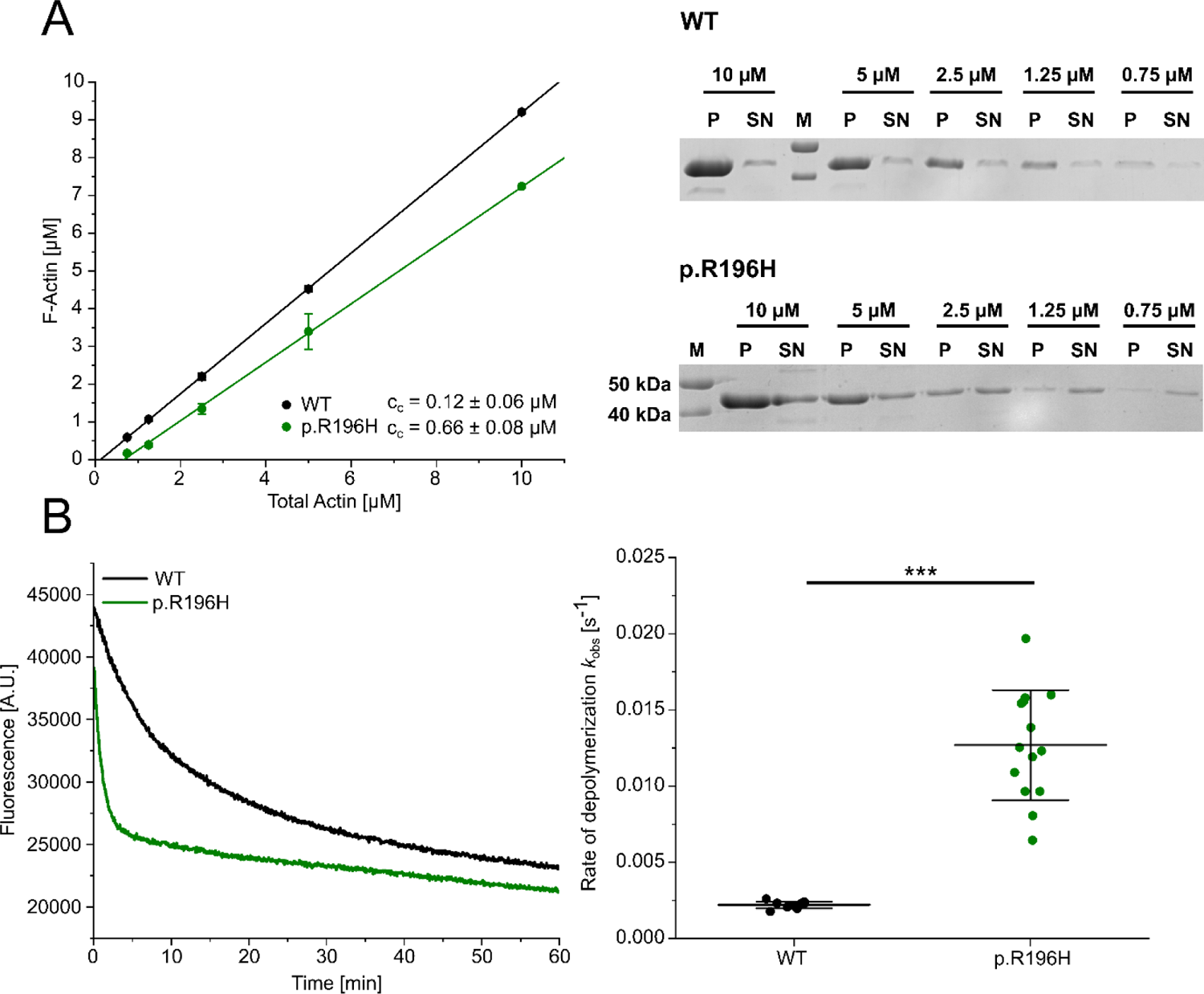
Determination of the critical concentration for polymerization and the apparent rate of filament depolymerization for p.R196H and WT β–actin. **(A)** High–speed sedimentation experiments with varying actin concentrations were used to determine the critical concentration for polymerization (c_c_) of p.R196H and WT actin. **(Left)** The amount of F–actin in the pellet fraction was determined by densitometry after SDS–PAGE. A linear regression was applied to the data, which yields the apparent critical concentration (c_c_) as the intercept with the x–axis. Data is shown as the mean ± SD of at least 3 individual experiments with the exception of p.R196H (10 µM), which is the result of one individual experiment. **(Right**) Exemplary SDS–gel of sedimentation experiments performed with p.R196H and WT actin. M: Molecular weight marker. **(B)** Pyrene–based dilution induced depolymerization experiments (5 % pyrene–labeled actin) were used to determine the apparent rate of filament depolymerization of p.R196H and WT filaments. The depolymerization of filaments was induced by the addition of G–buffer. **(Left)** Exemplary traces of individual depolymerization experiments performed with p.R196H and WT actin. **(Right)** Secondary plot summarizing the rates of filament depolymerization determined from all experiments. Every data point corresponds to an individual depolymerization experiment (N = 12 for WT, N = 14 for p.R196H). Data is shown as the mean ± SD.

To address the effect of the R196H mutation on spontaneous filament disassembly, we performed dilution–induced depolymerization experiments using pyrene–labeled pre–polymerized WT and p.R196H actin (**Figure 4B**). In accordance with a reduced filament stability, we observed a 5.8–fold increased depolymerization rate in experiments performed with p.R196H filaments compared to WT filaments.

### Arp2/3–mediated generation of branched actin networks is less efficient for p.R196H

The formation of branched actin structures absolutely requires the interaction of the actin filament with the Arp2/3 complex, one of the main regulators of actin nucleation *in vivo* (31). The Arp2/3 subunits Arp2 and Arp3 show high structural homology to actin and form with subunits ArpC1, ArpC2, ArpC3, ArpC4, and ArpC5 a stable seven–subunits complex that, after activation by a nucleation–promoting factor (NPF), associates with the side of an existing filament and serves as a nucleation core for the addition of new actin protomers (32–34). The newly added protomers intimately interact with Arp2 and Arp3, which form a stable dimer inside the complex–structure (35). Inspection of the high–resolution structure of the actin branch junction reveals that residue R196 is located in regions important for the interaction of Arp2 with the first protomer of the newly formed daughter filament (**Figure 5A**).

**Figure 5:**
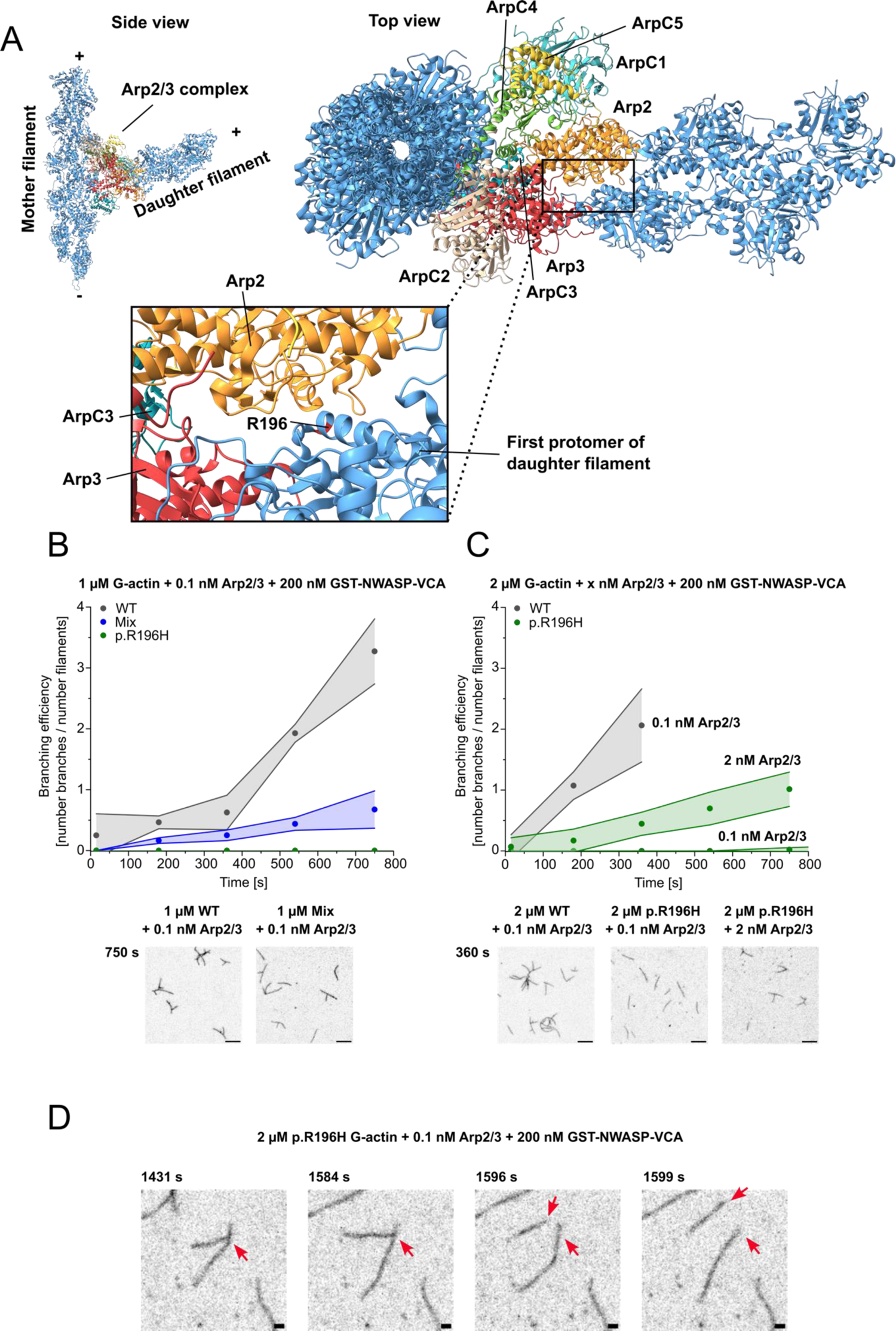
Analysis of the interaction of p.R196H and WT filaments with the Arp2/3 complex by TIRF microscopy. **(A)** Structural model showing the actin–Arp2/3 branch junction complex (based on PDB–ID: 7TPT (35)). The seven subunits of the Arp2/3 complex are differentially colored. The zoom– in shows the location of actin residue R196 at the interface between the initial protomer of the daughter filament and Arp2 of the Arp2/3 complex. **(B, C)** The formation of branched actin networks from p.R196H β–actin, WT β–actin, and a 1:1 mixture (10% ATTO–655 labeled) after salt-induced polymerization in the presence of human Arp2/3 complex (+GST–NWASP–VCA) was investigated by TIRF microscopy. The branching efficiency (number of Arp2/3–generated branches per initially linear filament) were determined for p.R196H and WT filaments at different actin and Arp2/3–complex concentrations. Plots show the determined branching efficiency over time. Data is shown as the mean ± SD. Exemplary micrographs show Arp2/3–generated branched actin structures at the indicated time after start of the experiment. Scale bar corresponds to 10 µm. (D) Time–lapse showing the spontaneous debranching (red arrows) of a p.R196H branched actin filament at the indicated experimental conditions. Scale bar corresponds to 1 µm.

We analysed the interaction between p.R196H filaments and activated human Arp2/3 complex using TIRFM. When used in single–filament TIRFM studies with 1 µM WT actin, 0.1 nM GST–N–WASP– VCA–activated human Arp2/3 complex was sufficient to induce the formation of branched actin structures (**Figure 5B**). As already observed in our polymerization studies, 1 µM p.R196H actin is not sufficient to generate enough linear filaments for branching efficiency studies. Therefore, we performed experiments with 1 µM of a 1:1 mix of p.R196H and WT actin. We observed enough initially linear filaments to quantify the branching efficiency. We quantified the branching efficiency of Arp2/3 complex by counting the number of branching points divided by the number of observed initially linear filaments at different time points. Comparing the branching efficiency to the branching efficiency in experiments performed with 1 µM WT, revealed a reduced efficiency for the heterofilaments, showing that the presence of p.R196H actin in the filament affects interaction with the Arp2/3 complex.

We increased the actin concentration to 2 µM p.R196H or WT actin to generate enough initial linear filaments in p.R196H samples to visualize the interaction of Arp2/3 complex with the pure mutant filaments (**Figure 5C**). We found almost no branching in experiments performed with p.R196H and 0.1 nM Arp2/3 complex. It is interesting to note that in experiments with p.R196H filaments we sometimes observed spontaneous debranching, a phenomenon that we never observed in experiments with WT actin (**Figure 5D**). Increasing the Arp2/3 complex concentration 20–fold in experiments with p.R196H still resulted in fewer observed branches compared to experiments with 2 µM WT actin and 0.1 nM Arp2/3 complex (**Figure 5C**).

Since the TIRFM–based assay does not allow an accurate quantitative assessment of branching efficiency under these conditions, we investigated the effect of higher Arp2/3 complex concentrations on WT and p.R196H actin in bulk polymerization studies using pyrene–labeled actin (**Figure 6**). Based on our previous results, we used 2 µM actin in our experiments to observe significant nucleation and elongation of p.R196H actin. The interpretation of experimental data from pyrene–based bulk– polymerization studies is based on the hypothesis that the two phases of actin polymerization, nucleation and elongation, give rise to two distinct observable phases in the temporal change of pyrenyl– fluorescence intensity. An initial phase in which pyrenyl–fluorescence increases only slowly (“lag– phase”), which is thought to be dominated by the initial nucleation of actin filaments, and a second phase in which pyrenyl–fluorescence increases more rapidly, which is interpreted as rapid barbed–end elongation. We determined the duration of the lag–phase (*t*_lag_) and the bulk–polymerization rate (*k*_obs_) by analyzing the two phases in experiments performed with 2 µM p.R196H or WT actin. We found an 11.5– fold increase of *t*_lag_ in experiments performed with p.R196H (*t*_lag_=917.5 ± 46.0 s) compared to experiments with WT actin (*t*_lag_=80.0 ± 10.6 s) (**Figure 6D**), while *k*_obs_ of p.R196H was 2.5–fold reduced compared to WT actin (**Figure 6C**). The addition of VCA–activated human Arp2/3 complex stimulated the polymerization of both p.R196H and WT actin and affected both observed phases, as previously described (33). The highest Arp2/3–concentration (40 nM) increased the *k*_obs_ of WT actin 109–fold and reduced the *t*_lag_ by 12–fold. In contrast, the highest Arp2/3–concentration had a milder effect on *k*_obs_ in experiments with p.R196H, resulting in only a 40–fold increase in *k*_obs_. Accordingly, the effect on *t*_lag_ was also less pronounced compared to WT experiments (6.5–fold decrease). As stated, we observed a pronounced increase in *t*_lag_ in experiments performed with p.R196H compared to WT actin in pyrene– based bulk experiments, while we observed only minimal differences in the nucleation efficiency between p.R196H actin and WT actin in our TIRFM–based experiments performed at the same actin concentration (2 µM).

**Figure 6:**
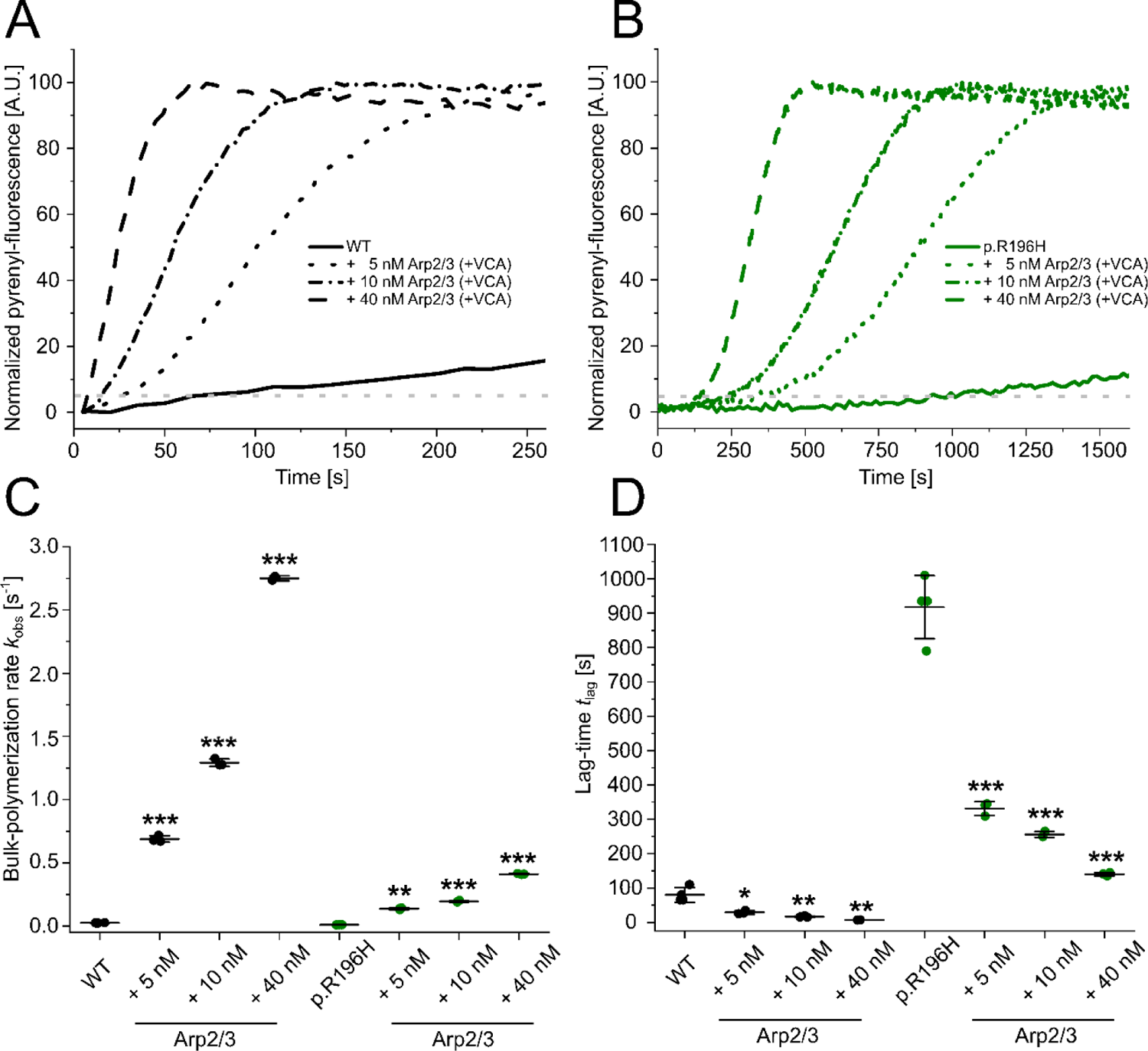
Analysis of the polymerization capacity of p.R196H and WT β–actin in the absence and presence of human Arp2/3 complex using bulk–polymerization experiments. **(A, B)** Pyrene–based polymerization experiment of 2 µM p.R196H or WT actin (5% pyrene–labeled) in the absence and presence of increasing concentrations of Arp2/3 (+ 200 nM GST–NWASP VCA). Shown are representative traces. Note the different x–axis in (A) and (B). The dotted grey lines indicate 5 % of the final pyrenyl–fluorescence, the threshold that was used to determine the lag–time *t*_lag_ in (D) **(C)** Bulk– polymerization rates determined from the experiments shown in (A) and (B). Every data point corresponds to an individual polymerization experiment (N = 3–4), as shown in (A) and (B). Data is shown as the mean ± SD. Individual values are summarized in Table 1. **(D)** Lag–time (*t*_lag_) determined from pyrene–based polymerization experiments. *t*_lag_ was defined as the time point when 5% of the final pyrenyl–fluorescence was reached, as indicated in (A) and (B) by the dotted grey line. Every data point corresponds to an individual polymerization experiment (N = 3–4), as shown in (A) and (B). Data is shown as the mean ± SD. Individual values are summarized in Table 1.

### Mutation R196H differentially affects interaction of F–actin with myosin isoforms

Residue R196 is buried inside the actin filament close to the filament axis and is therefore not directly involved in forming the actin–myosin interface. A previous study showed that a mutation in the hinge– region of the actin protomer affects actomyosin interaction in a myosin–isoform specific fashion, although the affected residue is buried inside the filament (36). This effect was linked to mutation– induced conformational changes of the actin filament that were maintained even in the presence of the filament–stabilizer phalloidin. To address the possible effect of mutation–induced conformational changes of the actin filament on the actomyosin complex, we analyzed the interaction between p.R196H and WT filaments and two distinctly different myosin isoforms using the unloaded *in vitro* motility assay. We chose human myosin–5A (Myo5A) and human non–muscle myosin–2A (NM2A) to reconstitute physiological relevant actomyosin complexes (**Figure 7**). TRITC–phalloidin labeled cytoskeletal β–actin filaments were moved by Myo5A with an average velocity of 352.2 ± 18.0 nm s^-1^. In contrast, filaments formed by p.R196H were moved 1.3–fold faster resulting in an average sliding velocity of 457.1 ± 38.6 nm s^-1^. We observed no significant change in the velocity with which p.R196H and WT filaments were moved by NM2A, indicating a similar interaction with the stress fiber associated myosin (**Figure 7**, **Table 1**).

**Figure 7:**
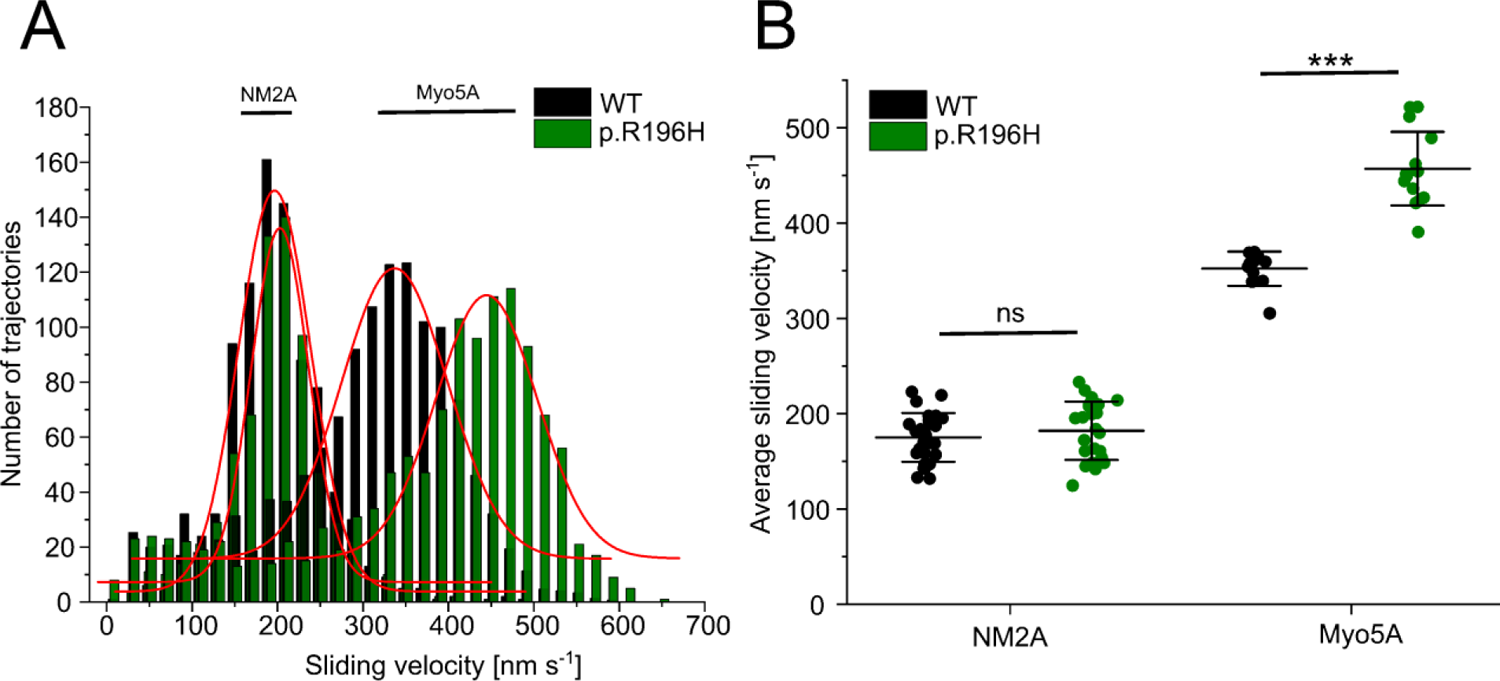
Analysis of the interaction of human NM2A and Myo5A with p.R196H and WT β–actin filaments. **(A)** The interaction of surface–immobilized NM2A–HMM and Myo5A–HMM with TRITC– phalloidin stabilized WT β–actin and p.R196H filaments was probed using the unloaded *in vitro* motility assay. Shown are representative velocity distributions obtained from recorded trajectories of WT and p.R196H filaments in a single experiment. A Gaussian fit (red line) was applied to the obtained velocity distributions in each experiment to determine the average sliding velocity of the respective filaments. **(B)** Secondary plot showing all measured sliding velocities. Each data point represents a single experiment (N=27 for WT filaments on NM2A, N=23 for p.R196H filaments on NM2A, N=12 for WT filaments on Myo5A, N=15 for p.R196H filaments on Myo5A). The average sliding velocity in each experiment was determined as described in (A). The mean ± SD of all performed experiments is shown.

## Discussion

Baraitser–Winter cerebrofrontofacial syndrome (BWCFF) is the best–defined human syndrome associated with cytoskeletal β–actin variants, and patients present with a relatively homogeneous phenotype (7,8,10,15). However, the link between the molecular perturbations of cytoskeletal actin dynamics caused by the mutated actin molecule and the phenotype of the patients remains largely unknown. *In vitro* characterization of the observed mutation–induced defects remains a fundamental task in understanding the genotype–phenotype correlation, as it allows dissection of individual steps of actin dynamics in the absence and in the presence of ABPs. Variant p.R196H is the most frequently observed variant causing BWCFF. Affected patients show prototypical craniofacial anomalies and neurodevelopmental disorders, which are thought to be related to neuronal migration defects (7,8), making this variant particularly interesting for the study of the molecular mechanisms of actinopathies.

Our study provides evidence that rapid protein degradation of p.R196H leading to functional haploinsufficiency is unlikely to be a contributing disease mechanism, as p.R196H monomers are properly folded, stable and have nucleotide interactions comparable to WT β–actin protein. Using TIRF microscopy–based single–filament studies with fluorescently labeled actin, we have shown that mutation R196H results in a reduced rate of barbed–end elongation and a reduced rate of filament nucleation at low actin concentrations (< 2 µM). Both observed defects are attenuated in the presence of equimolar concentrations of WT actin, mimicking the expected outcome of heterozygosity in patients. Our observations suggest a disease mechanism involving the stable production of p.R196H protomers and their efficient incorporation into actin filaments, thereby perturbing cytoskeletal filament dynamics and their interactions with ABPs. Differential scanning fluorimetry and pyrene–actin based bulk depolymerization experiments of p.R196H and WT filaments provide further support for this interpretation, revealing the inherent instability of p.R196H filaments and a compromised G– to F–actin ratio.

A previous study used a cell–free *in vitro* translation system to produce the variant p.R196C and showed that it folds as efficiently as WT protein (37). Furthermore, the authors presented data that indicate a polymerization defect for the p.R196C variant, as filaments formed by the p.R196C variant *in vitro* are significantly shorter than WT filaments under the same conditions. These data suggest a comparable polymerization defect for the p.R196C variant and the p.R196H variant characterized in our study. It is therefore reasonable to assume that all BWCFF–associated variants at position R196 disrupt the monomer–filament equilibrium by perturbing important intra–protomer interactions that stabilize the protomer region close to the filament axis. Furthermore, due to the close proximity of the affected residue to the interstrand contact formed between the neighboring residue E195 and K113 of the opposite protomer, weakening of the interstrand contacts may contribute to the observed reduced filament stability (**Figure 1B**).

R196H is located in SD3 of the actin protomer, adjacent to the helical axis of the actin filament, well away from the canonical interfaces for G– and F–actin binding proteins (22). However, this is not the case with the Arp2/3 complex, where the Arp2 and Arp3 subunits of the complex mimic a nucleation core for a daughter filament on the side of a mother filament. Therefore, we hypothesized that mutations that affect actin–actin interactions, as is the case for the p.R196H variant, impair the interaction of the variant protomers with the Arp2 and Arp3 subunits, reducing the efficiency of branch formation. R196 is located in close proximity to the Arp2 subunit in the branch junction complex (**Figure 5A**). Consistent with this view, we observe a reduced efficiency of branch formation and an apparent branch instability using TIRFM–based single–filament studies and pyrene–actin bulk–polymerization experiments. Previously, perturbed actin–Arp2/3 interaction were reported in studies investigating the molecular consequences of variants p.K118N/M in cytoskeletal γ–actin causing non–syndromic hearing loss (38,39). In the context of BWCFF, neuronal migration defects are suggested to be causative for the occurrence of pachygyria in the patients carrying p.R196H leading to developmental disabilities (7,10). The Arp2/3 complex is an essential component of the protein machinery that drives directed cell migration (31) and is important for human corticogenesis (40,41). Several steps in human corticogenesis are directly dependent on intact Arp2/3 complex function, including the morphogenesis and migration of radial glial and oligodendrocyte precursors (41–43) and the migration of the mature neuron (40). Furthermore, mutations in a repressor of the Arp2/3 complex, α–catenin, lead to pachygyria, thereby providing a link between misregulated Arp2/3 complex function and cortical malformations (44). This suggests that perturbed interactions between variant p.R196H and the Arp2/3 complex contribute to the phenotype of the patients.

Various myosin isoforms were shown to be essential for neuronal function. Myo5A is highly abundant in the brain and its importance for neuronal function is well established (45). Various Myo5A–mediated transport processes contribute to synaptic plasticity (45), motor learning (46) and neurotransmitter recycling (47,48). NM2A is enriched in the vasculature of the brain, where it contributes to angiogenesis (49). We observed an effect of the p.R196H variant on the interaction of Myo5A with the actin filament, as exemplified by a 1.3–fold increase in filament sliding velocity on surface–immobilized Myo5A– HMM, whereas the variant did not affect the velocity of filaments moving on surface–immobilized NM2A–HMM. Therefore, isoform–specific perturbances of the actomyosin interactions potentially contribute to the cortical malformations and neurodevelopmental disorders of the patients.

## Material and Methods

### Plasmids

The coding sequence of human cytoskeletal β–actin (Uniprot–ID: P60709) was fused via a C–terminal linker (ASR(GGS)_3_A) to a His_8_–tagged thymosin–β4 moiety (Uniprot–ID: P62328) and was cloned into the multiple cloning site of the pFastBac–Dual vector under control of the polyhedrin promotor. The plasmid encoding the p.R196H variant was generated by site–specific mutagenesis of the WT plasmid and verified by sequencing. The pFastBac1 vectors for the production of the HMM–like constructs of human NM2A and human Myo5A were described previously (Hundt et al, 2016; Reindl et al, 2022). The coding sequences of human myosin light chains MYL6a (Uniprot–ID: P60660) and MYL12b (Uniprot–ID: O14950) were cloned into the pFastBac–Dual–vector (Hundt et al, 2016; Reindl et al, 2022). Human calmodulin (Uniprot–ID: P0DP23) was cloned into the pFastBac–Dual vector. The human Arp2/3 complex consisting of subunits Arp2 (Uniprot–ID: P61160), Arp3 (Uniprot–ID: P61158), ArpC1B (Uniprot–ID: O15143), ArpC2 (Uniprot–ID: O15144), ArpC3A (Uniprot–ID: O15145), ArpC4 (Uniprot–ID: P59998), ArpC5 (Uniprot–ID: O15511) and a C–terminal FLAG tag on ArpC3A was produced using the expression vector pBIG2abc, provided by Roberto Dominguez (University of Pennsylvania, Philadelphia, USA). The VCA domain of human N–WASP (Uniprot–ID: O00401) was cloned into pGEX–6P–2 vector. The cDNA encoding human thymosin–β4 (Uniprot–ID: P62328) was a gift from Taro Uyeda (Waseda University, Tokyo, Japan) and subcloned into the pET23(+) vector to generate a His_8_–tagged construct. The generation of recombinant bacmids and viruses for the production of proteins in the Sf9 system followed the Bac–to–Bac baculovirus expression system protocol (Thermo Fisher Scientific, Waltham, MA, USA).

### Protein production and purification

Recombinant human cytoskeletal β–actin WT and the mutant p.R196H were produced in the form of actin–thymosin–β4 fusion protein in the baculovirus/*S. frugiperda* (Sf9) insect cell expression system and purified following a protocol previously described for human cytoskeletal γ–actin (25). Myosin HMM–like constructs were co–produced with human Myl6/Myl12b (NM2A–HMM) or human calmodulin (Myo5A–HMM) in Sf9 cells and purified as described previously (50,51). Production and purification of the human Arp2/3 complex followed the published protocol by Zimmet *et al.* (52). The human GST–N–WASP–VCA construct was produced in ArcticExpress (DE3) cells and purified as previously described (25).

Human thymosin–β4–His_8_ was produced in Rosetta2 cells. For a typical purification, cells from 1 liter of expression culture were resuspended in 100 mL of lysis buffer (10 mM TRIS pH 7.8, 100 mM NaCl, 15 mM imidazole, 5 mM β–mercaptoethanol, 1 mM PMSF, 100 µg/mL TAME, 80 µg/mL TPCK, 2 µg/mL pepstatin, 5 µg/mL leupeptin) supplemented with 250 µg/mL lysozyme (from hen egg white; Merck KGaA, Darmstadt, Germany) and incubated on ice for 30 minutes. The cells were lysed by sonification and treated with DNase–I (Roche, Basel, Switzerland) for 30 minutes on ice to remove bacterial DNA. The lysate was cleared by centrifugation at 35,000 × g for 30 minutes and the cleared supernatant was subsequently added to 3 mL of Pure Cube^TM^ NiNTA material (Cube Biotech, Monheim am Rhein, Germany) equilibrated with lysis buffer. The suspension was incubated for 2 hours at 4 °C under constant rotation. The beads were washed with a 25–fold excess of wash buffer (10 mM TRIS pH 7.8, 50 mM NaCl, 15 mM imidazole) and loaded onto a gravity–flow column. The protein was eluted with 250 mM imidazole pH 7.5 in 3 mL fractions. The fractions containing the pure protein were pooled and diluted to a final concentration of 10 mM imidazole pH 7.5. The diluted sample was concentrated using a Vivaspin 20 concentrator (MWCO: 5,000; Sartorius, Göttingen, Germany) to a concentration of at least 1 mM. The concentration was determined by SDS–gel electrophoresis followed by densitometry at a ChemiDoc–MP–gel documentation system (Bio–Rad Laboratories, Inc., Hercules, USA) due to the absence of aromatic amino acids in thymosin–β4. A fragment of the C2–domain of myosin–binding– protein C (MyBPC) of known size (17 kDa) and concentration was used as a standard. The final protein was snap–frozen in liquid nitrogen and stored at –80 °C until used.

### DNase–I inhibition assay, DSF and nucleotide exchange assay

DNase–I inhibition assays were performed as previously described (17,25). Differential scanning fluorimetry (DSF) experiments with monomeric and filamentous WT and mutant actin followed protocols previously described (17,25), with slight modifications. Briefly, for DSF experiments with G–actin in the presence of thymosin–β4 the G–actin/thymosin–β4 complex was formed by incubating equimolar amounts of both proteins on ice for 30 minutes prior to measurements. For DSF experiments with phalloidin–stabilized actin filaments, filaments were incubated overnight at 4 °C with a 1.5–fold molar excess of phalloidin. The thermal denaturation temperature T_M_ was determined from the peak of the first derivative of the melting curve. The full width at half maximum (FWHM) correlates with the cooperativity of the observed melting process.

The rate of nucleotide dissociation for monomeric p.R196H and β–actin WT was determined using ε– ATP (Jena Bioscience, Jena, Germany) at a HiTech Scientific SF61 stopped–flow system (TgK Scientific Limited, Bradford on Avon, UK), as previously described (25).

### TIRF microscopy–based assays

Filament nucleation and elongation of WT and p.R196H actin were determined in TIRF microscopy– based assays using freshly clarified protein. WT and p.R196H actin were labeled at cysteine–375 with ATTO–655 (ATTO–TEC, Siegen, Germany) to visualize nucleation and elongation of actin filaments, as previously described (17,25). The glass coverslips used for the assembly of flow cells were cleaned and surfaces were chemically treated following previously described protocols (17,25). Actin polymerization was induced by diluting the G–actin solution to a final concentration of 0.5 µM, 1 µM or 2 µM (10% labeled actin) with 2× TIRF–buffer (20 mM imidazole, 50 mM KCl, 1 mM MgCl_2_, 1 mM EGTA, 0.2 mM ATP, 15 mM glucose, 20 mM β–ME, 0.25 % methylcellulose, 0.1 mg/mL glucose oxidase and 0.02 mg/mL catalase). After mixing, the solutions were immediately flushed into flow cells and image acquisition was started. For experiments performed in the presence of VCA–activated human Arp2/3 complex, VCA–activated Arp2/3 complex was prediluted in KMEI buffer (10 mM imidazole pH 7.4, 50 mM KCl, 1 mM MgCl_2_, 1 mM EGTA) and diluted to the final concentration in TIRF–buffer before addition of actin.

Image series were acquired at an Olympus IX83 inverted fluorescence microscopes (Olympus, Hamburg, Germany) equipped with a 60×/1.49 NA PlanApo TIRF oil immersion objective and an Orca Flash 4.0 CMOS camera (Hamamatsu Photonics Deutschland GmbH, Herrsching, Germany). Image analysis was performed using ImageJ (53). The elongation rates of individual actin filaments were determined by manually tracking individual filaments. To determine the branching efficiency in the presence of Arp2/3, the number of branching points were manually counted at several time points and divided by the number of initially linear filaments.

### Sedimentation experiments

Actin was polymerized for 3 hours at room temperature by diluting G–actin into F–buffer (10 mM TRIS pH 7.8, 100 mM KCl, 5 mM MgCl_2_, 0.5 mM EGTA, 0.1 mM DTT, 0.1 mM ATP) in 1.5 mL polypropylene centrifuge tubes (Beckman Coulter, Brea, CA, USA) suited for high–speed centrifugation to final concentrations ranging from 0.75 µM – 10 µM. Samples were centrifuged at 136,000 × *g* (30 min, 4°C) and the resulting pellet and supernatant fraction were subjected to SDS–PAGE. The gels were imaged at a ChemiDoc–MP–gel documentation system (Bio–Rad Laboratories, Hercule, CA, USA). The amount of actin in the pellet and the supernatant was determined by densitometry in the Image Lab software (Bio–Rad Laboratories, Hercule, CA, USA). The concentration of F–actin in each sample was plotted against the total actin concentration in each sample. A linear regression was fitted to the complete dataset, which yields the critical concentration of actin polymerization (c_c_) as the intercept with the x–axis.

### Pyrene–actin based bulk–polymerization assays

The pyrene–actin based bulk–polymerization experiments in the absence and presence of VCA– activated Arp2/3 complex were performed at a Synergy 4™ microplate reader (BioTek Instruments, Winooski, VT, USA) using the built–in filter set (Excitation: 340/30 nm, Emission: 400/30 nm) at 25 °C, as previously described (25). For dilution–induced depolymerization experiments, 20 µM of pre– polymerized pyrene–labeled F–actin was rapidly diluted to 0.1 µM by addition of G–buffer (10 mM TRIS pH 7.8, 0.2 mM CaCl_2_, 0.2 mM ATP).

Bulk–polymerization rates were determined according to Doolittle *et al.* (54) by fitting a linear regression to the linear region around the time–point of half–maximal fluorescence. The lag–time (*t*_lag_) of the reaction was defined as the time–point at which the reaction reaches 5 % of the final fluorescence signal. The rate of filament disassembly determined from dilution–induced depolymerization experiments was calculated by fitting a mono–exponential function to the obtained data, including a linear component that accounts for bleaching. Labeling of WT and p.R196H actin at cysteine–374 using *N*–(1–pyrene)iodoacetamide (Thermo Fisher Scientific, Waltham, MA, USA) followed established protocols (**55**).

### Unloaded *in vitro* motility assay

Unloaded *in vitro* motility assays to analyze the interaction between WT and p.R196H actin filaments and NM2A–HMM or Myo5A–HMM were performed as previously described (17,50). Flow cells for the use in the *in vitro* motility assay were constructed by using nitrocellulose–coated coverslips. Prior to the experiments, purified NM2A–HMM was incubated with the GST–tagged kinase domain (residues 1425–1776) of human smooth muscle myosin light chain kinase (Uniprot–ID: A0A8I5KU53) at a molar ratio of 10:1 in 25 mM MOPS pH 7.3, 50 mM KCl, 5 mM MgCl_2_, 1 mM CaCl_2_, 0.2 µM calmodulin, 3 µM regulatory light chain (MYL12b), 3 µM essential light chain (MYL6a), 1 mM DTT and 1 mM ATP for 30 minutes at 30 °C. This is necessary, as phosphorylation of the MYL12b is absolutely required for full activity of NM2A–HMM. Myo5–HMM was incubated with 2 µM human calmodulin and each buffer was supplemented with calmodulin to prevent calmodulin dissociation from Myo5A–HMM during the experiments. An Olympus IX70 inverted fluorescence microscopes (Olympus, Hamburg, Germany) equipped with a 60×/1.49 NA PlanApo oil immersion objective and an Orca Flash 4.0 CMOS camera (Hamamatsu Photonics Deutschland GmbH, Herrsching, Germany) was used for time–lapse image acquisition. Experiments were performed at a 37°C. The ImageJ Plugin wrmTrck (56) was used to determine the trajectories and corresponding velocities of the individual actin filaments.

### In silico analysis

Assessment and visualization of the potential implications of variant p.R196H for actin structure was performed using ChimeraX (57)

## Data analysis

Data analysis and graph plotting were performed with Origin 2023 (OriginLab Corporation. Massachusetts, USA). Errors are given as standard deviation (SD) based on three independent experiments if not otherwise specified. The significance of the data was evaluated in Origin 2023 using a two–sample t–test (p > 0.05 ≙ ns, p ≤ 0.05 ≙ *, p ≤ 0.01 ≙ **, p ≤ 0.001 ≙ ***, p ≤ 0.0001 ≙ ****).

## Author Contributions

J.N.G. purified proteins; J.N.G. performed experiments; J.N.G. and D.J.M. analyzed the data; J.N.G designed figures; J.N.G. and D.J.M. conceived and coordinated the study and wrote the manuscript; D.J.M. was responsible for funding acquisition, supervision and project administration.

## Acknowledgments

The authors thank Roberto Dominguez (University of Pennsylvania, Philadelphia, USA) for providing vector pBIGBac–Arp2/3 and Taro Uyeda (Waseda University, Tokyo, Japan) for providing human thymosin–β4 cDNA. D.J.M was supported by grants from Deutsche Forschungsgemeinschaft (DFG) (MA1081/23–1, MA1081/28–1), J.N.G. and D.J.M. are members of the European Union’s Horizon 2020 research and innovation program under the EJP RD COFUND–EJP N° 825575 with support from the German Federal Ministry of Education and Research under Grant Agreement 01GM1922B. D.J.M. is a member of the Cluster of Excellence RESIST (EXC 2155; DFG–Project ID: 39087428–B11). J.N.G. is supported by the HiLF I grant for early career researcher from Medizinische Hochschule Hannover. We gratefully acknowledge support provided by the Core Unit for Structural Biochemistry and the Core Unit for Laser Microscopy at MHH.

## Materials Availability

Unique reagents generated in this study are available from the lead contacts with a completed Materials Transfer Agreement.

## Literature

1. Perrin BJ, Ervasti JM. The actin gene family: function follows isoform. Cytoskelet Hoboken NJ. 2010 Oct;67(10):630–4.

2. Ponti A, Machacek M, Gupton SL, Waterman-Storer CM, Danuser G. Two Distinct Actin Networks Drive the Protrusion of Migrating Cells. Science. 2004 Sep 17;305(5691):1782– 6.

3. Pollard TD, Blanchoin L, Mullins RD. Molecular Mechanisms Controlling Actin Filament Dynamics in Nonmuscle Cells. Annu Rev Biophys Biomol Struct. 2002;

4. Ridley AJ, Schwartz MA, Burridge K, Firtel RA, Ginsberg MH, Borisy G, et al. Cell Migration: Integrating Signals from Front to Back. Science. 2003 Dec 5;302(5651):1704–9.

5. Plastino J, Blanchoin L. Dynamic stability of the actin ecosystem. J Cell Sci. 2019;132(4).

6. Pollard TD. Actin and actin-binding proteins. Cold Spring Harb Perspect Biol. 2016;8(8).

7. Verloes A, Di Donato N, Masliah-Planchon J, Jongmans M, Abdul-Raman OA, Albrecht B, et al. Baraitser-Winter cerebrofrontofacial syndrome: Delineation of the spectrum in 42 cases. Eur J Hum Genet. 2015;23(3):292–301.

8. Rivière JB, Van Bon BWM, Hoischen A, Kholmanskikh SS, O’Roak BJ, Gilissen C, et al. De novo mutations in the actin genes ACTB and ACTG1 cause Baraitser-Winter syndrome. Nat Genet. 2012;44(4):440–4.

9. Latham SL, Ehmke N, Reinke PYA, Taft MH, Eicke D, Reindl T, et al. Variants in exons 5 and 6 of ACTB cause syndromic thrombocytopenia. Nat Commun. 2018 Dec;9(1).

10. Di Donato N, Rump A, Koenig R, Der Kaloustian VM, Halal F, Sonntag K, et al. Severe forms of Baraitser-Winter syndrome are caused by ACTB mutations rather than ACTG1 mutations. Eur J Hum Genet. 2014;22(2):179–83.

11. Cuvertino S, Stuart HM, Chandler KE, Roberts NA, Armstrong R, Bernardini L, et al. ACTB Loss-of-Function Mutations Result in a Pleiotropic Developmental Disorder. Am J Hum Genet. 2017;

12. Wijk E, Krieger E, Kemperman MH, De Leenheer EMR, Huygen PLM, Cremers CWRJ, et al. A mutation in the gamma actin 1 (ACTG1) gene causes autosomal dominant hearing loss (DFNA20/26). J Med Genet. 2003 Dec;40(12):879–84.

13. Zhu M, Yang T, Wei S, DeWan AT, Morell RJ, Elfenbein JL, et al. Mutations in the γ-Actin Gene (ACTG1) Are Associated with Dominant Progressive Deafness (DFNA20/26). Am J Hum Genet. 2003;73(5):1082–91.

14. Rendtorff NDND, Zhu M, Fagerheim T, Antal TL, Jones MP, Teslovich TM, et al. A novel missense mutation in ACTG1 causes dominant deafness in a Norwegian DFNA20/26 family, but ACTG1 mutations are not frequent among families with hereditary hearing impairment. Eur J Hum Genet. 2006;

15. Baraitser M, Winter RM. Iris coloboma, ptosis, hypertelorism, and mental retardation: a new syndrome. J Med Genet. 1988 Jan 1;25(1):41–3.

16. Costa CF, Rommelaere H, Waterschoot D, Sethi KK, Nowak KJ, Laing NG, et al. Myopathy mutations in alpha-skeletal-muscle actin cause a range of molecular defects. J Cell Sci. 2004 Jul 1;117(Pt 15):3367–77.

17. Greve JN, Schwäbe FV, Pokrant T, Faix J, Di Donato N, Taft MH, et al. Frameshift mutation S368fs in the gene encoding cytoskeletal β-actin leads to ACTB-associated syndromic thrombocytopenia by impairing actin dynamics. Eur J Cell Biol. 2022 Apr 1;101(2):151216.

18. Hundt N, Preller M, Swolski O, Ang AM, Mannherz HG, Manstein DJ, et al. Molecular mechanisms of disease-related human β-actin mutations p.R183W and p.E364K. FEBS J. 2014;281(23):5279–91.

19. Müller M, Mazur AJ, Behrmann E, Diensthuber RP, Radke MB, Qu Z, et al. Functional characterization of the human α-cardiac actin mutations Y166C and M305L involved in hypertrophic cardiomyopathy. Cell Mol Life Sci. 2012;69(20).

20. Bryan KE, Wen KK, Zhu M, Rendtorff ND, Feldkamp M, Tranebjaerg L, et al. Effects of human deafness gamma-actin mutations (DFNA20/26) on actin function. J Biol Chem. 2006 Jul 21;281(29):20129–39.

21. Schröder JM, Durling H, Laing N. Actin myopathy with nemaline bodies, intranuclear rods, and a heterozygous mutation in ACTA1 (Asp154Asn). Acta Neuropathol (Berl). 2004 Sep;108(3):250–6.

22. Dominguez R, Holmes KC. Actin Structure and Function. Annu Rev Biophys. 2011 Jun 9;40(1):169–86.

23. Noguchi TQP, Kanzaki N, Ueno H, Hirose K, Uyeda TQP. A novel system for expressing toxic actin mutants in Dictyostelium and purification and characterization of a dominant lethal yeast actin mutant. J Biol Chem. 2007 Sep 21;282(38):27721–7.

24. Hatano T, Sivashanmugam L, Suchenko A, Hussain H, Balasubramanian MK. Pick-ya actin – a method to purify actin isoforms with bespoke key post-translational modifications. J Cell Sci. 2020 Jan 30;133(2):jcs241406.

25. Greve JN, Marquardt A, Heiringhoff R, Reindl T, Thiel C, Di Donato N, et al. The non-muscle actinopathy-associated mutation E334Q in cytoskeletal γ-actin perturbs interaction of actin filaments with myosin and ADF/cofilin family proteins. Sellers JR, Dötsch V, editors. eLife. 2024 Mar 6;12:RP93013.

26. Schüler H, Lindberg U, Schutt CE, Karlsson R. Thermal unfolding of G-actin monitored with the DNase I-inhibition assay. Stabilities of actin isoforms. Eur J Biochem. 2000;

27. Lo MC, Aulabaugh A, Jin G, Cowling R, Bard J, Malamas M, et al. Evaluation of fluorescence-based thermal shift assays for hit identification in drug discovery. Anal Biochem. 2004 Sep 1;332(1):153–9.

28. Dedova IV, Nikolaeva OP, Safer D, De La Cruz EM, dos Remedios CG. Thymosin β4 Induces a Conformational Change in Actin Monomers. Biophys J. 2006 Feb 1;90(3):985–92.

29. Levitsky DI, Pivovarova AV, Mikhailova VV, Nikolaeva OP. Thermal unfolding and aggregation of actin. FEBS J. 2008;275(17):4280–95.

30. Das S, Ge P, Oztug Durer ZA, Grintsevich EE, Zhou ZH, Reisler E. D-loop Dynamics and Near-Atomic-Resolution Cryo-EM Structure of Phalloidin-Bound F-Actin. Structure. 2020;28(5).

31. Goley ED, Welch MD. The ARP2/3 complex: an actin nucleator comes of age. Nat Rev Mol Cell Biol. 2006 Oct;7(10):713–26.

32. Beltzner CC, Pollard TD. Pathway of Actin Filament Branch Formation by Arp2/3 Complex*. J Biol Chem. 2008 Mar 14;283(11):7135–44.

33. Mullins RD, Heuser JA, Pollard TD. The interaction of Arp2/3 complex with actin: Nucleation, high affinity pointed end capping, and formation of branching networks of filaments. Proc Natl Acad Sci. 1998;

34. Pollard TD, Beltzner CC. Structure and function of the Arp2/3 complex. Curr Opin Struct Biol. 2002;

35. Ding B, Narvaez-Ortiz HY, Singh Y, Hocky GM, Chowdhury S, Nolen BJ. Structure of Arp2/3 complex at a branched actin filament junction resolved by single-particle cryo-electron microscopy. Proc Natl Acad Sci. 2022 May 31;119(22):e2202723119.

36. Noguchi TQP, Komori T, Umeki N, Demizu N, Ito K, Iwane AH, et al. G146V mutation at the hinge region of actin reveals a myosin class-specific requirement of actin conformations for motility. J Biol Chem. 2012;287(29).

37. Machida K, Miyawaki S, Kanzawa K, Hakushi T, Nakai T, Imataka H. An *in Vitro* Reconstitution System Defines the Defective Step in the Biogenesis of Mutated β-Actin Proteins. ACS Synth Biol. 2021 Nov 19;10(11):3158–66.

38. Jepsen L, Kruth KA, Rubenstein PA, Sept D. Two Deafness-Causing Actin Mutations (DFNA20/26) Have Allosteric Effects on the Actin Structure. Biophys J. 2016;111(2).

39. Kruth KA, Rubenstein PA. Two deafness-causing (DFNA20/26) actin mutations affect Arp2/3-dependent actin regulation. J Biol Chem. 2012 Aug 3;287(32):27217–26.

40. Chou FS, Wang PS. The Arp2/3 complex is essential at multiple stages of neural development. Neurogenesis. 2016 Dec 27;3(1):e1261653.

41. Wang PS, Chou FS, Ramachandran S, Xia S, Chen HY, Guo F, et al. Crucial roles of the Arp2/3 complex during mammalian corticogenesis. Development. 2016 Aug 1;143(15):2741–52.

42. Mair DB, Elmasli C, Kim JH, Barreto AD, Ding S, Gu L, et al. The Arp2/3 complex enhances cell migration on elastic substrates. Mol Biol Cell. 2023 Jun;34(7):ar67.

43. Li Y, Wang PS, Lucas G, Li R, Yao L. ARP2/3 complex is required for directional migration of neural stem cell-derived oligodendrocyte precursors in electric fields. Stem Cell Res Ther. 2015 Mar 21;6(1):41.

44. Schaffer AE, Breuss MW, Caglayan AO, Al-Sanaa N, Al-Abdulwahed HY, Kaymakçalan H, et al. Biallelic loss of human CTNNA2, encoding αN-catenin, leads to ARP2/3 complex overactivity and disordered cortical neuronal migration. Nat Genet. 2018 Aug;50(8):1093– 101.

45. Hammer JA, Wagner W. Functions of Class V Myosins in Neurons. J Biol Chem. 2013 Oct 4;288(40):28428–34.

46. Wagner W, Brenowitz SD, Hammer JA. Myosin-Va transports the endoplasmic reticulum into the dendritic spines of Purkinje neurons. Nat Cell Biol. 2011 Jan;13(1):40–8.

47. Röder IV, Petersen Y, Choi KR, Witzemann V, Hammer JA, Rudolf R. Role of Myosin Va in the plasticity of the vertebrate neuromuscular junction in vivo. PloS One. 2008;3(12):e3871.

48. Röder IV, Choi KR, Reischl M, Petersen Y, Diefenbacher ME, Zaccolo M, et al. Myosin Va cooperates with PKA RIα to mediate maintenance of the endplate in vivo. Proc Natl Acad Sci. 2010 Feb 2;107(5):2031–6.

49. Ma X, Uchida Y, Wei T, Liu C, Adams RH, Kubota Y, et al. Nonmuscle myosin 2 regulates cortical stability during sprouting angiogenesis. Mol Biol Cell. 2020 Aug 15;31(18):1974– 87.

50. Reindl T, Giese S, Greve JN, Reinke PY, Chizhov I, Latham SL, et al. Distinct actin– tropomyosin cofilament populations drive the functional diversification of cytoskeletal myosin motor complexes. iScience. 2022 Jul 15;25(7):104484.

51. Hundt N, Steffen W, Pathan-Chhatbar S, Taft MH, Manstein DJ. Load-dependent modulation of non-muscle myosin-2A function by tropomyosin 4.2. Sci Rep. 2016 Feb 5;6:20554.

52. Zimmet A, Van Eeuwen T, Boczkowska M, Rebowski G, Murakami K, Dominguez R. Cryo-EM structure of NPF-bound human Arp2/3 complex and activation mechanism. Sci Adv. 2020 Jun;6(23):eaaz7651.

53. Rueden CT, Schindelin J, Hiner MC, DeZonia BE, Walter AE, Arena ET, et al. ImageJ2: ImageJ for the next generation of scientific image data. BMC Bioinformatics. 2017 Nov 29;18(1):529.

54. Doolittle LK, Rosen MK, Padrick SB. Measurement and analysis of in vitro actin polymerization. Methods Mol Biol. 2013;1046.

55. Cooper JA, Walker SB, Pollard TD. Pyrene actin: documentation of the validity of a sensitive assay for actin polymerization. J Muscle Res Cell Motil. 1983 Apr;4(2):253–62.

56. Nussbaum-Krammer CI, Neto MF, Brielmann RM, Pedersen JS, Morimoto RI. Investigating the spreading and toxicity of prion-like proteins using the metazoan model organism C. elegans. J Vis Exp JoVE. 2015 Jan 8;(95):52321.

57. Pettersen EF, Goddard TD, Huang CC, Meng EC, Couch GS, Croll TI, et al. UCSF ChimeraX: Structure visualization for researchers, educators, and developers. Protein Sci Publ Protein Soc. 2021 Jan;30(1):70–82.

58. Schutt CE, Myslik JC, Rozycki MD, Goonesekere NCW, Lindberg U. The structure of crystalline profilin-β-actin. Nature. 1993;

59. Arora AS, Huang HL, Singh R, Narui Y, Suchenko A, Hatano T, et al. Structural insights into actin isoforms. Lappalainen P, Akhmanova A, Galkin V, editors. eLife. 2023 Feb 15;12:e82015.

